# Structural insights into PFAS-β-lactoglobulin binding mechanism mediating PFAS toxicity

**DOI:** 10.1101/2025.09.02.673760

**Authors:** Shalja Verma, Anika Singh, Randhal S. Ramirez Orozco, Lela Vukovic, Mahesh Narayan, Pravindra Kumar

## Abstract

The strong, polar-covalent nature of C-F bonds contributes to the forever nature of per- and polyfluoroalkyl (PFAS) substances. PFAS are toxic to humans. Here, we have examined the ability of the small, globular, milk protein β-lactoglobulin to bind PFAS. The protein transports hydrophobic and amphiphilic compounds, including retinol and fatty acids, for vision and brain development; therefore, underscoring its interactions with PFAS is significant. The crystal structures of β-lactoglobulin complexed with PFOA (Perfluorooctanoic acid) at 2.0 (Å), PFOS (Perfluorooctanesulfonic acid) at 2.5 (Å), and PFDA (Perfluorodecanoic acid) at 2.0 (Å) reveal high affinity of the compounds for the central calyx of β-lactoglobulin, which is the canonical retinol and fatty acid binding site. Analyses of the data indicate significant hydrophobic interactions stabilizing the binding of the PFAS hydrophobic “tails” within the calyx and interactions between Lys60 and Lys69 and PFAS polar head groups. Comparative structural analysis revealed the presence of an open conformation of the EF loop containing the Glu89 latch residue in the complexed structures vis-a-vis the apo-form. Molecular dynamics (MD) simulations revealed high stability of the PFAS binding and attainment of energy minima in all complexes. The average binding energy of PFDA in β-lactoglobulin calyx was -25 kcal/mol, which was higher than PFOS (-21 kcal/mol) and PFOA (-23 kcal/mol) due to increased van der Waals interactions of the longer hydrophobic chain of PFDA with β-lactoglobulin. This work advances a mechanism by which β-lactoglobulin can recruit PFAS and act as a transporter for the “forever” chemical, potentially mediating its neurotoxicity.

## Introduction

The C-F bond is very stable due to the strong electronegative character of the fluorine atom (4.0 versus 2.5 for carbon), resulting in the shared electron density essentially becoming concentrated around the fluorine and leaving the carbon relatively electron poor. As a consequence, C-F bonds possess significant polarity (dipole moment) and an electrostatic character. The partial charges (C^δ+^—F^δ−^) on the fluorine and carbon are attractive and contribute to its unusual bond strength.^1–3^ For example, C-F linkages can possess bond dissociation energies up to 130 kcal/mol, which makes the bond strength of C–F higher than other carbon–halogen and carbon–hydrogen bonds. Lastly, the three lone pairs in the C-F bonded fluorine atoms coupled with the aforementioned partial negative charge create “shields” which are both steric shield and electrostatic in nature. The “shield” effectively diminishes the ability of the bonded carbon atom to nucleophilic attack. Per- and polyfluoroalkyl (PFAS) chemicals contain C-F bonds and are therefore termed as “forever” by nature. The described C-F bond characteristics are contributors to PFAS inertness.^4–5^

Another unique aspect of C-F bonds, which impact the interaction of PFAS molecules in biological systems, is the weak/absent hydrogen bond donor capability of the F in the said linkage.^6^ In classical hydrogen-bond-forming species such as water or HF, the electronegative oxygen atom (or fluorine atom) makes the H atom a hydrogen bond donor with the lone pairs of the oxygen (or fluorine) acting as acceptors. However, PFAS molecules are not readily capable of making hydrogen bonds through the lone pairs in their fluorinated “tails”. This is because the strongly electronegative nature of the fluorine prevents the molecule’s lone pairs from being available to a hydrogen donor. The low polarization of the s and p electrons in F also makes it a poor hydrogen bond acceptor despite the high electronegativity or lone pairs. Thus, though the carbon-fluorine (C-F) bond is very polar, its fluorine is a very weak hydrogen bond acceptor.^4–5^

The lack of a strong ability to make intra- and intermolecular interactions by the C-F chain in PFAS, arising from the low polarizability of fluorine, makes for more volatile compounds with lower boiling points when compared to counterparts of similar molecular masses. Importantly, the entropic penalty paid for introducing PFAS into water can therefore only be compensated for by the “hydrophobic effect” and makes for the “oleophobic” effect of PFAS.^6^

How such physico-chemical tendencies influence the interactions between C-F rich PFAS molecules and biological assemblies is of interest considering their presence in human organs, tissues, biological fluids and other aqueous bio-matrices. Furthermore, interpreting the conversations between PFAS and biological systems is also of clinical significance considering their association with adverse human health outcomes which constitutes major societal and governmental concerns.^7^ Human exposure to PFAS can occur through cookware, ingestion of contaminated drinking water, food, and food packaging. Ingestion can occur through the use of some PFAS-containing consumer products (lipstick, upholstery, hygiene products, diapers, waterproof clothing, stain-resistant carpets and fabrics, and cleaning products), the inhalation of contaminated dust or airborne particles, or dermal contact with contaminated materials.^8^

PFAS exposure is linked to obesity, decreased fertility, developmental anomalies in foetuses, neonates, and children, compromised immunity, and increased risk of prostate, kidney, and testicular cancers.^9–12^ Studies have shown that PFAS can cross the blood-brain barrier and may contribute to neuroinflammation, neurotoxicity, cognitive and locomotory decline, all of which promote the progression of Alzheimer’s (AD) and Parkinson’s diseases (PD).^13–17^ A recent comprehensive study focused on how they affect gene expression of neuronal-like cells.^18^ Exposure to a suite of C-F bond-containing organic moieties resulted in modest but distinct changes in lipids and caused over 700 genes to express differently, including those involved in synaptic growth and neural function. The findings also hint that the neurotoxic effects are structure-dependent, highlighting the chemical complexities involved (C-F chain-length, head-group, and branching).^18^ Elsewhere, a study revealed that multigenerational exposure of the nematode *C. elegans* to perfluoronoanoic acid (PFNA) significantly increased mutation frequencies of progeny by preferentially inducing T: A → C: G substitutions and small insertion-deletions within repetitive regions.^19^ Mechanistic analyses suggested that PFNA elicited DNA double-strand breaks.

Of concern is the detection of PFAS in serum from the umbilical cord and newborn blood, demonstrating that these chemicals are capable of passing through the placental barrier with estimates of the efficiency of placental transfer ranging between 30-79%.^20^ Elsewhere, breast milk has been shown to account for 83-99% of PFAS total daily intake for infants.^21–22^ A study involving 50 lactating mothers revealed 16 PFAS compounds in 4-100% of the samples.^23^ Therein, perfluorodecanoic acid (PFDA) was found in 94% of the cohort, with the median residing at 7.4 pg/ml. Perfluorooctanesulfonic acid (PFOS) and perfluorooctanoic acid (PFOA) were the most abundant PFAS in these samples (medians 30.4 and 13.9 pg/mL, respectively). PFAS presence has also been detected in infant formulas either innately or post-reconstitution and in bovine milk.^24–26^

The hydrophobic nature of PFAS is implicated in its ability to bind to blood serum proteins (albumin and globulins).^27–28^ The presence of proteins in milk, including the hydrophobic ligand transport protein β-lactoglobulin, provides the motivation to examine the impact of PFAS exposure on the structure and function of milk proteins.

β-lactoglobulin is a major protein of ruminant milk that accounts for 10-15 % of its protein content.^29–30^ It contributes to a major fraction of whey protein and acts as a transporter for hydrophobic metabolites and amphiphilic compounds, regulates the activity of pregastic lipase in the neonatal stage, and protects retinol from light-mediated degradation.^31^ From the aforementioned literature, it is highly likely that contaminated milk matrices provide a conduit for the interaction of β-lactoglobulin and PFAS. Using biochemical, biophysical, and in-silico approaches, previous studies have proposed a high binding affinity of PFAS for β-lactoglobulin and including its potential for the transport of PFAS to different organs.^32–35^ However, structural studies revealing the exact binding mechanism of the PFAS in β-lactoglobulin utilizing X-ray crystallography have not been conducted. Such studies can be beneficial for devising the binding and transport mechanism of PFAS by β-lactoglobulin and identifying binding inhibition approaches to reduce milk contaminants.

In this study, we determined the crystal structures of β-lactoglobulin in complex with PFOA, PFOS, and PFDA, revealing structural intricacies mediating the binding of PFAS in the β-lactoglobulin central calyx. Furthermore, in-depth molecular dynamics simulation analysis of the crystal structures was performed to analyze the stability and dynamic changes communicated by the binding of PFAS in β-lactoglobulin.

The analyses form part of our overarching objective to develop a precise understanding of the electrostatic, van der Waals, H-bonding, London Dispersion forces, along with hydrophobic effects, further modulated by the nature of the C-F moieties (chain-length, head-group, branching), that may potentially impact human health. This study begins to lay the roadmap for mechanisms by which commonly used and environmentally prevalent C-F containing chemistries corrupt protein structures and interfere with their physiological function. It provides the molecular basis by which PFAS can compromise nutritional sufficiency in infants, children, and adults and act as a transporter for the “forever” chemical, potentially driving pathological outcomes.

## Materials and methods

### Co-Crystallization of β-lactoglobulin-PFAS complex

Bovine β-lactoglobulin, PFDA, PFOA, and PFOS were procured from Sigma Aldrich. The lyophilized protein was dissolved in MiliQ water up to a concentration of 80 mg/ml and delipidated using activated charcoal at pH 3. The concentration of the delipidated protein was obtained to be approximately 40 mg/ml.^36^ The protein was incubated with the PFAS compounds for 10 min at 20 °C at a concentration of 10 mM. Crystallization was performed using the sitting drop method at a protein to reservoir ratio of 1:3, and screening of different crystallization conditions was performed using crystallization screens to obtain complex crystals of β-lactoglobulin with PFAS.^37^ The cube-shaped crystals of β-lactoglobulin in complex with PFAS were obtained in a crystallization condition containing 1.8 M sodium phosphate monobasic monohydrate and potassium phosphate dibasic at pH 8.2.

### Data Collection and structure solving

The collection of X-ray data was performed using a Rigaku Micromax-007 HF microfocus high-intensity rotating anode X-ray generator installed with a Rigaku Hypix 6000C detector available at home source at the Macromolecular Crystallography Unit, IIC, IIT Roorkee. The diffraction data indexing and integration were performed using CrysAlisPro Software. The data scaling and merging were done using the AIMLESS program of the CCP4i2 v.8.0.019 suite, followed by initial phasing by molecular replacement, considering the apo structure of β-lactoglobulin (PDB ID: 1B0O) as the search model, using the MOLREP program of the CCP4i2 suite.^28–40^ Iterative rounds of model building were performed using the COOT program, and the refinement of the structure at each step was performed using the REFMAC program of the CCP4i2 suite.^41–42^ The difference Fourier maps of the PFAS-β-lactoglobulin complexes revealed electron densities of PFAS compounds at levels above 3σ, permitting modelling of PFAS into the density. Omit maps were made using Phenix v.1.18.2-3874.^43^ The 3D and 2D interaction images were generated using ChimeraX 1.8 and PyMol v.2.5.5.^44–45^

### Molecular Dynamics Simulation

Molecular dynamic simulations of crystal structures of β-lactoglobulin-PFAS complexes and apo β-lactoglobulin were performed using GROMACS 2019.5 software^46^ for a 100 ns duration in two independent runs for each system to study the dynamics of interactions between the protein and PFAS. The missing residues of flexible regions in crystal structures due to the absence of electron density were added using Coot software.^41^ The topologies of PFAS compounds were generated using CGenFF web server integrated with the CHARMM force field, while the protein topology was generated by applying the CHARMM36 force field using the pdb2gmx tool of GROMACS.^47–48^ The apo-protein and complexes were then solvated in a dodecahedron box. The solvated systems were neutralized by sodium ions and then were subjected to energy minimization at a maximum force limit of 10 kJ/mol. Further, number of atoms, volume, temperature (NVT), and number of atoms, pressure, temperature (NPT) equilibrations each for 100 ps were performed. Eventually, each system was subjected to a MD production run of 100 ns each. The trajectories of MD simulations were analyzed for Root Mean Square Deviations (RMSD), Root Mean Square Fluctuations (RMSF), Radius of gyration (Rg), Solvent Accessible Surface Area (SASA), B-factor, and secondary structure changes during the simulation.^49^ The PCA analysis of the trajectory frames/conformations was performed. The clustering analysis of the simulation trajectories was performed to evaluate the most consistent interactions during the simulation trajectories.^50^ The end state free binding energy analysis for the last 40ns of the simulation trajectories was performed using gmx_mmpbsa v1.6.4 in duplicate.^51^ The interaction images of the trajectories were generated using PyMol v.2.5.5, and the values of all the parameters were plotted using Origin 2024b software (https://www.originlab.com/).^45^

## Results and discussion

### Crystal structures of β-lactoglobulin in complex with PFAS compounds

The crystal structures of β-lactoglobulin in complex with PFOS, PFOA, and PFDA were determined in the P 3_2_ 2 1 space group with one molecule in an asymmetric unit. Statistics pertaining to data collection and refinement of three complex structures are presented in Table 1. The 162 amino acid long β-lactoglobulin protein (MW∼ 18,400) folds into 8 anti-parallel β-strands forming the β-barrel or the calyx, a characteristic of lipocalin family proteins. The 9^th^ strand, lying near the first strand, is responsible for dimerization, and a 3-turn alpha-helix lies at the outer surface (Figure 1A, B). Two intrachain disulfide bonds, Cys60-Cys160 and Cys106-Cys119, were observed, and the density for the residues Ala111-Glu114 in the GH loop was partial or missing in all three complex structures due to the high flexibility of this loop.^36^ The EF loop, which acts as a gate over the calyx, was observed in an open conformation, and the side chain of the latch residue Glu89 was shifted outward from the calyx to enable binding of the PFAS compounds.^33–34^ Previous studies reported pH-dependent changes in the Glu89 side chain conformation, associating the exposed conformation of Glu89 with pH 8.2 and the buried conformation at pH 6.2 due to the “Tanford transition”. A similar open conformation of EF loop, along with the Glu89 residue, was observed in the PFAS-β-lactoglobulin complexes obtained in the current study at pH 8.2, indicating the contribution of the “Tanford transition” in mediating PFAS binding in the central calyx of β-lactoglobulin.^35^

**Figure 1:**
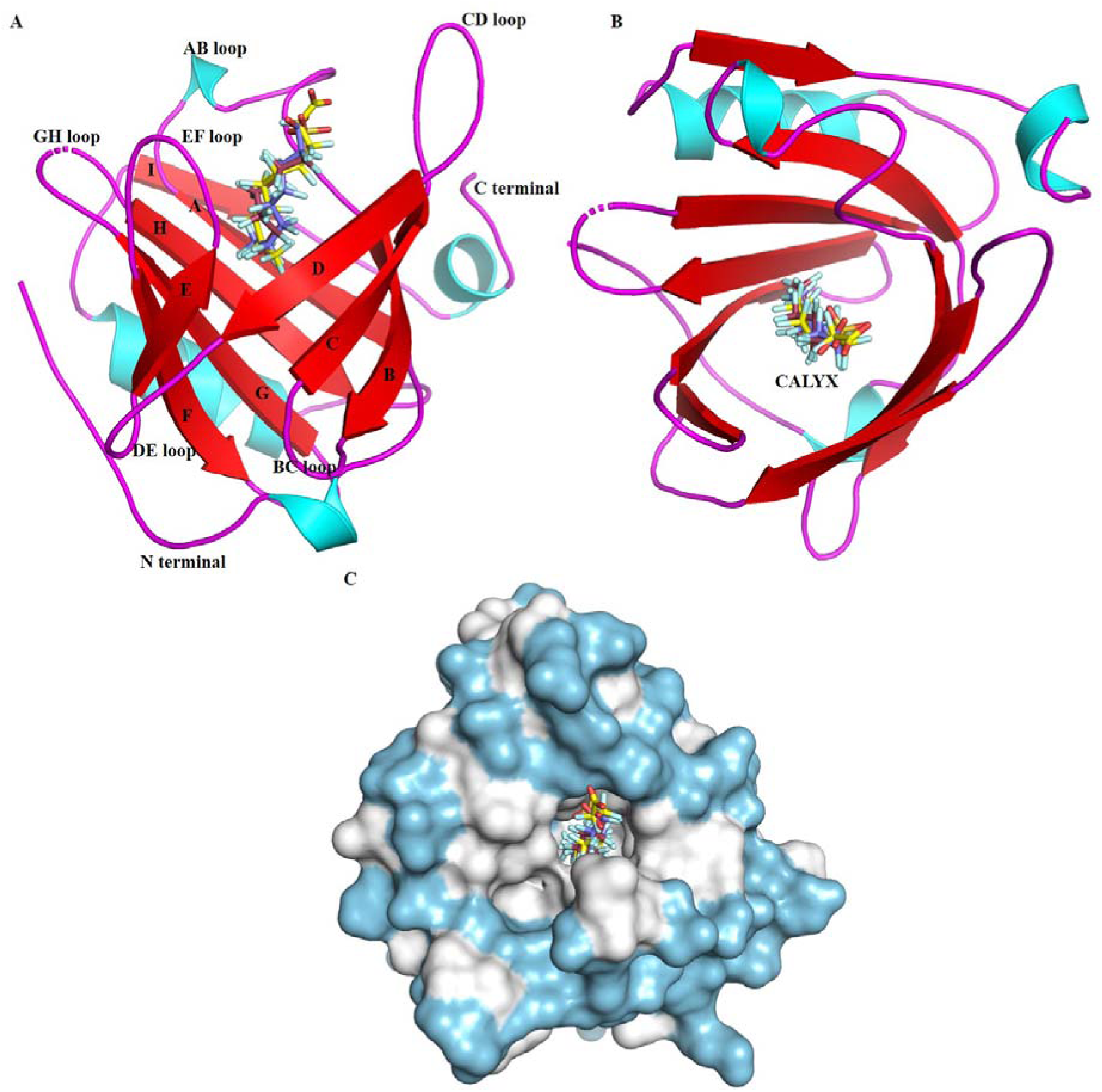
Superimposition of crystal structures of β-lactoglobulin in complex with PFOS (ruby), PFOA (blue), PFDA (yellow) A). Vertical view showing 9 beta-strands interspersed by loops and helices forming a central calyx with all three compounds bound into it. B) Horizontal view showing the same binding site inside the central calyx for all three compounds. C) Hydrophobic surface view of beta-lactoglobulin showing the presence of highly hydrophobic residues inside the central calyx. (Hydrophobic residue: white, hydrophilic residues: blue)

**Table 1:**
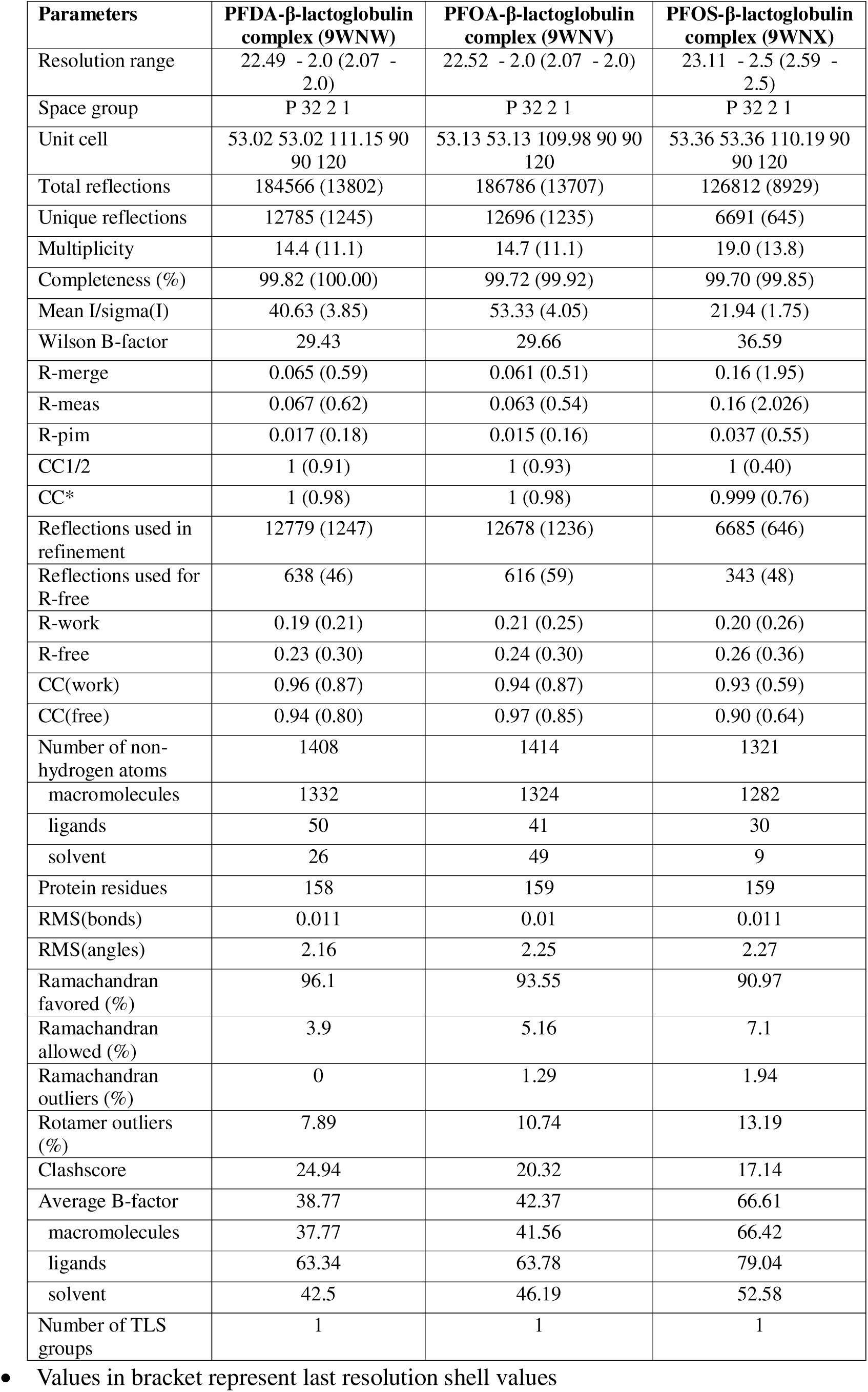
Refinement statistics of the crystal structures of PFDA-β-lactoglobulin, PFOA-β-lactoglobulin, and PFOS-β-lactoglobulin complexes.

The difference Fourier maps showing electron densities for the respective PFAS compounds were observed in the central cavity formed by the 8-β-strand calyx in complex structures (Figure 2). The hydrophobic “tail” of the PFAS compounds extends inside the hydrophobic cavity, forming hydrophobic interactions with Leu39, Ile56, Leu58, Ile71, Ile84, Val92, Phe105, Met107, and Val41, while the hydrophilic polar head group was stabilized by the hydrogen bond interactions with Lys60 and Lys69 at the cavity entrance (Figure 1C). The placement of the phenyl ring of Phe105 just in front of the terminal CF_3_ and subsequent CF_2_ groups of the PFAS compounds in all three crystal complex structures hindered the compounds from extending straight deep into the calyx cavity and led to a curved orientation of the compounds. Also, steric repulsion among fluorine atoms along with their large van der Waals radius, in PFAS compounds, causes twisting of the carbon backbone, resulting in 15/7 helix conformation, compared to the zig-zag conformation of the hydrocarbon analog, as observed in the crystal structures.^6^

**Figure 2:**
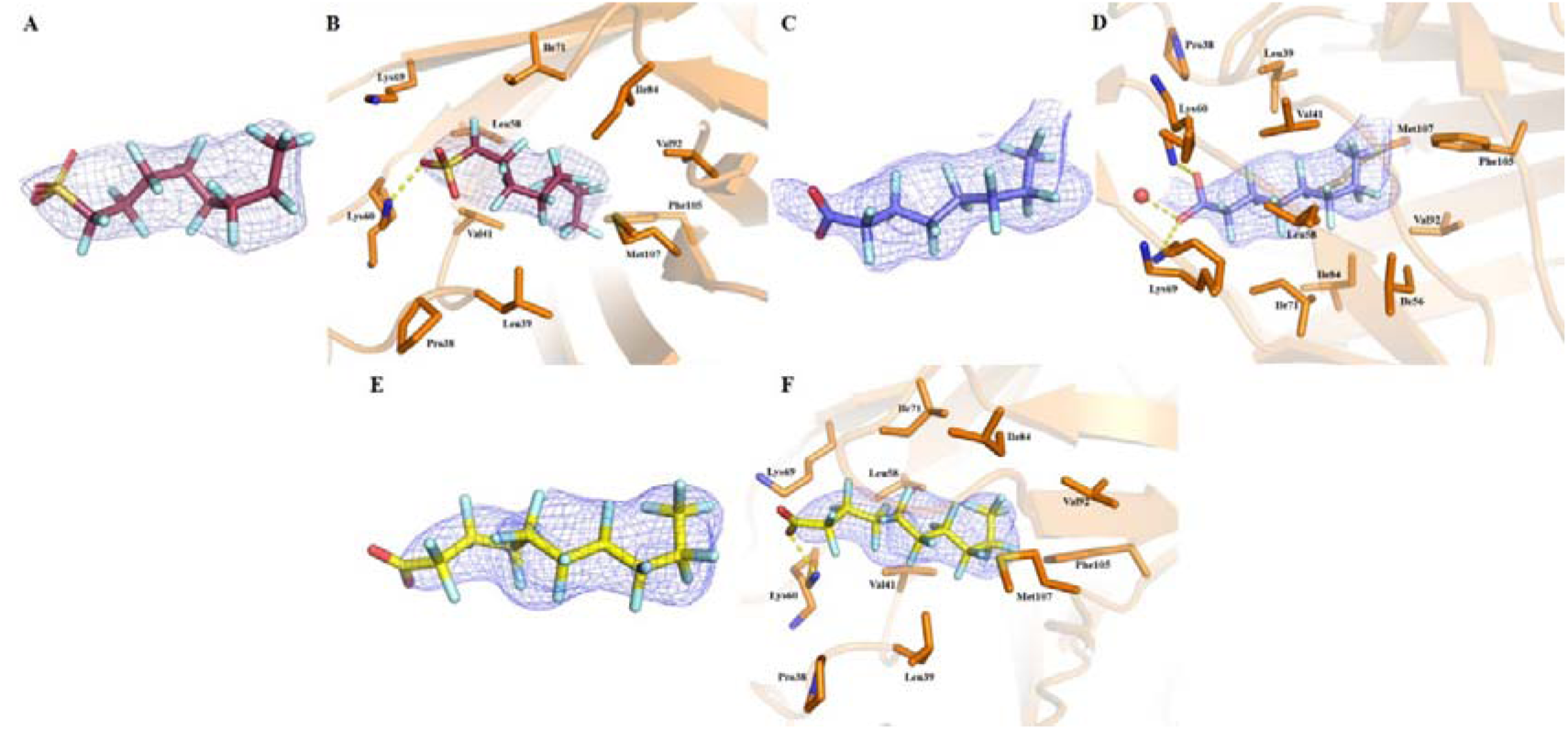
Crystal structures of β-lactoglobulin with PFAS. A) Electron density Fo-Fc map contoured at 3σ of PFOS in complex with β-lactoglobulin at 2.5 Å B) Stick representation of interacting residues (orange) with PFOS (ruby) in the central calyx of β-lactoglobulin. C) Electron density F_o_-F_c_ map contoured at 3σ of PFOA in complex with β-lactoglobulin at 2.0Å D) Stick representation of interacting residues (orange) with PFOA (blue) in the central calyx of beta-lactoglobulin E) Electron density Fo-Fc map contoured at 3σ of PFDA in complex with β-lactoglobulin at 2.0 Å F) Stick representation of interacting residues (orange) with PFDA (yellow) in the central calyx of β-lactoglobulin

The crystal structure of β-lactoglobulin-PFOS complex was solved at 2.5 Å resolution with *R_work_* of 0.21 and *R_free_* of 0.26. The side chains of Gln5, Lys60, and Ser150 residues were present in alternate conformations. The O3 oxygen atom of the sulphonic acid group formed a hydrogen bond with the NZ nitrogen atom of Lys60 at a distance of 2.84 Å, while the O2 oxygen atom of the sulphonic group was at a distance of 3.13 Å from the same nitrogen atom of Lys60. Further, the NZ nitrogen atom of Lys69 was at a hydrogen bond distance of 3.2 Å from the O3 oxygen of the sulphonic acid group of PFOS, revealing potential interactions. Similar interactions of the carboxyl group of palmitates at a distance of 2.72 Å and 3.22 Å with the NZ nitrogen atom of Lys60 and Lys69, respectively, have been observed in previous studies, stabilizing the binding of palmitate in the calyx of β-lactoglobulin (Figure 2A and B).^36^

In the β-lactoglobulin-PFOA complex at 2 Å resolution with *R_work_* and *R_free_* of 0.21 and 0.25, the side chain alternate conformations of Gln5, Ser30, Gln35, Lys47, Lys60, Lys69, Thr125, and Ile162 residues were present in alternate conformations. The O1 oxygen atom of the carboxyl group of the PFOA formed a hydrogen bond with the NZ nitrogen atom of Lys69 at a distance of 3.30 Å. A water molecule lying in close vicinity to the carboxyl group of PFOA, in turn, formed a hydrogen bond with the O1 oxygen atom of PFOA at a distance of 2.64 Å. The O2 oxygen atom of PFOA formed a hydrogen bond with the NZ nitrogen atom of Lys60 at a distance of 2.92 Å (Figure 2C and D).

The complex crystal structure of β-lactoglobulin-PFDA was determined at 2 Å resolution, and *R_free_* and *R_work_* values were 0.19 and 0.24, respectively, with Val3, Thr18, Asp28, Gln59, Lys60, Ile72, Leu95, Val118, Cys119, Lys135, Met145, and Glu158 having alternate side chain conformations. The PFDA bound to β-lactoglobulin exhibited hydrogen bonding of the NZ nitrogen atom of Lys60 at a distance of 2.49 Å with the O2 oxygen atom of the carboxyl group of PFDA, thus stabilizing the binding of PFDA in the calyx of β-lactoglobulin (Figure 2E and F). Previous structural studies have reported similar hydrogen bond interactions of Lys60 and Lys69 residues of β-lactoglobulin in stabilizing the binding of long-chain fatty acids having structural homology with PFAS compounds.^36, 52^

### Comparison of β-lactoglobulin PFAS complexes open conformation with apo-β-lactoglobulin closed conformation

Comparative analysis of β-lactoglobulin PFAS complexes at pH 8.2 with the apo β-lactoglobulin at 8.2 (PDB ID: 2BLG) revealed a few structural changes (Figure 3). The loop formed by residues Ile84-Glu89 was shifted towards the central calyx in β-lactoglobulin PFAS complexes compared to apo-form. Furthermore, residues Ile12-Val15 were found to be in helix conformation in β-lactoglobulin PFAS complexes in contrast with being in a loop conformation in apo-β-lactoglobulin. Also, the electron density of residues between Ser110-Glu114 forming loop conformation was absent in the β-lactoglobulin PFAS complexes compared to previously reported apo-form where all residues were modeled in the electron density. Lastly, analyses of average B factors across all residues with and without PFOA demonstrated differences as seen in Figure 3C. These differences may arise from a combination of PFAS-associated effects in addition to variations in crystallographic conditions across the two studies (this versus PDB ID:2BLG). These structural variations are indicative of allosteric effects of PFAS binding (Figure 3).

**Figure 3:**
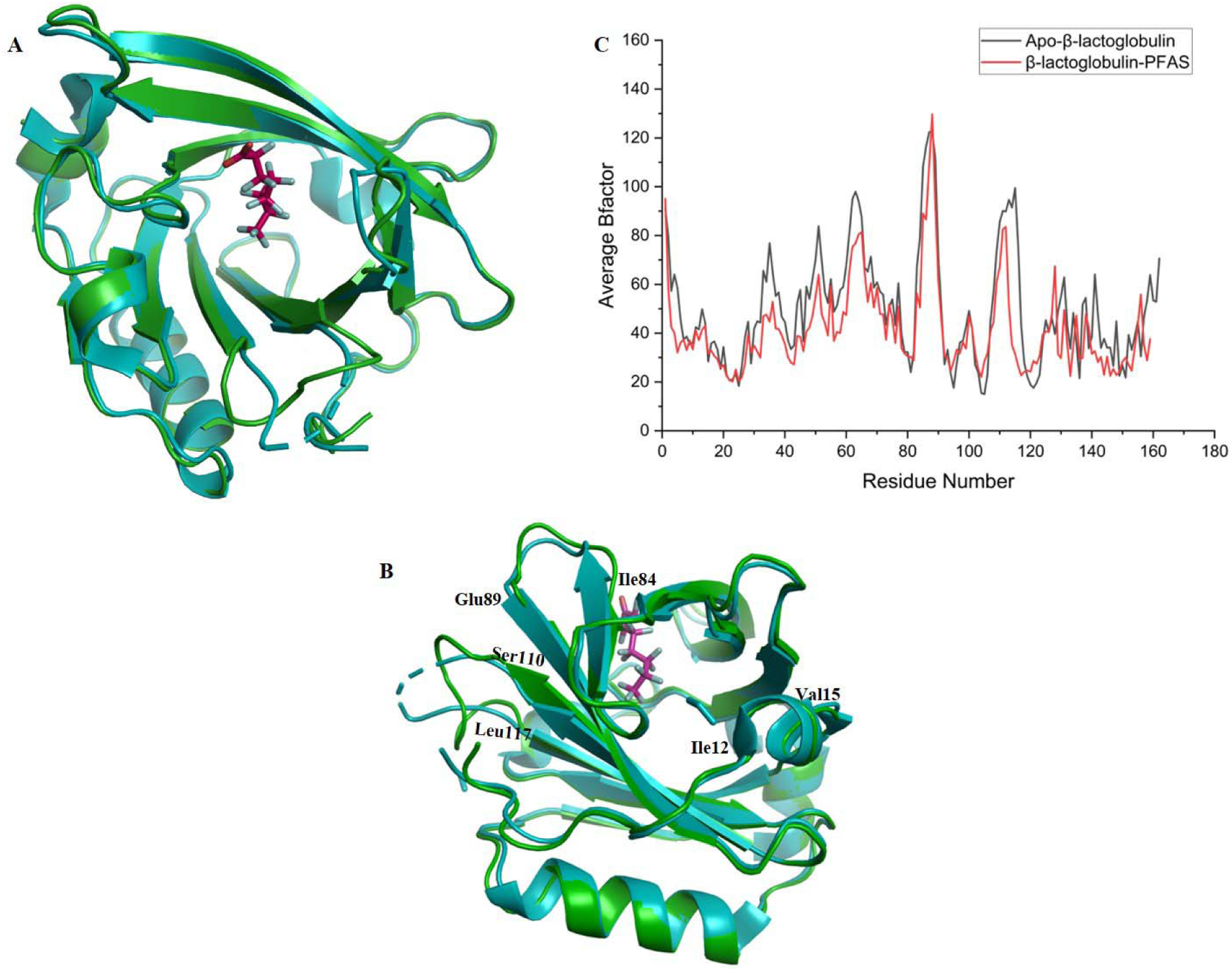
Comparative structural analysis of β-lactoglobulin PFOA complex (PDB ID: 9WNV, This study) and the apo β-lactoglobulin (PDB ID: 2BLG) at pH 8.2. A) Cartoon representation of β-lactoglobulin PFAS complex (cyan) superimposed over the apo β-lactoglobulin (green). B) Side view of superimposed cartoon representation of β-lactoglobulin PFOA complex (cyan) and the apo β-lactoglobulin (green) displaying structural changes. C) Average b-factor versus residue plot of β-lactoglobulin PFOA complex and the apo β-lactoglobulin.

Furthermore, structural superimposition of the β-lactoglobulin PFAS complexes crystallized at pH 8.2 over the apo-β-lactoglobulin form crystallized at pH 5.2 (PDB ID: 2AKQ previous study) and pH 6.2 (PDB ID: 3BLG, previous study) revealed an angular displacement of ∼ 64.6° in outward direction in the EF loop (at pH 8.2) in all PFAS complexes.^35, 53^. This conformational change in the EF loop is mediated by the variation in pH, where acidic pH leads to closure of the loop while neutral or basic pH results in open conformation of the loop (Figure 4A). The surface representation depicts complete closure of the central cavity by the EF loop and the side chain of Glu89, the latch residue, at acidic pH in the apo-β-lactoglobulin structures, while an open central cavity was visible in the PFAS-β-lactoglobulin complexes at pH 8.2 reported in this study as well as in apo-β-lactoglobulin at basic pH 8.2 in previous study (Figure 4B, C, D).^35, 53^

**Figure 4:**
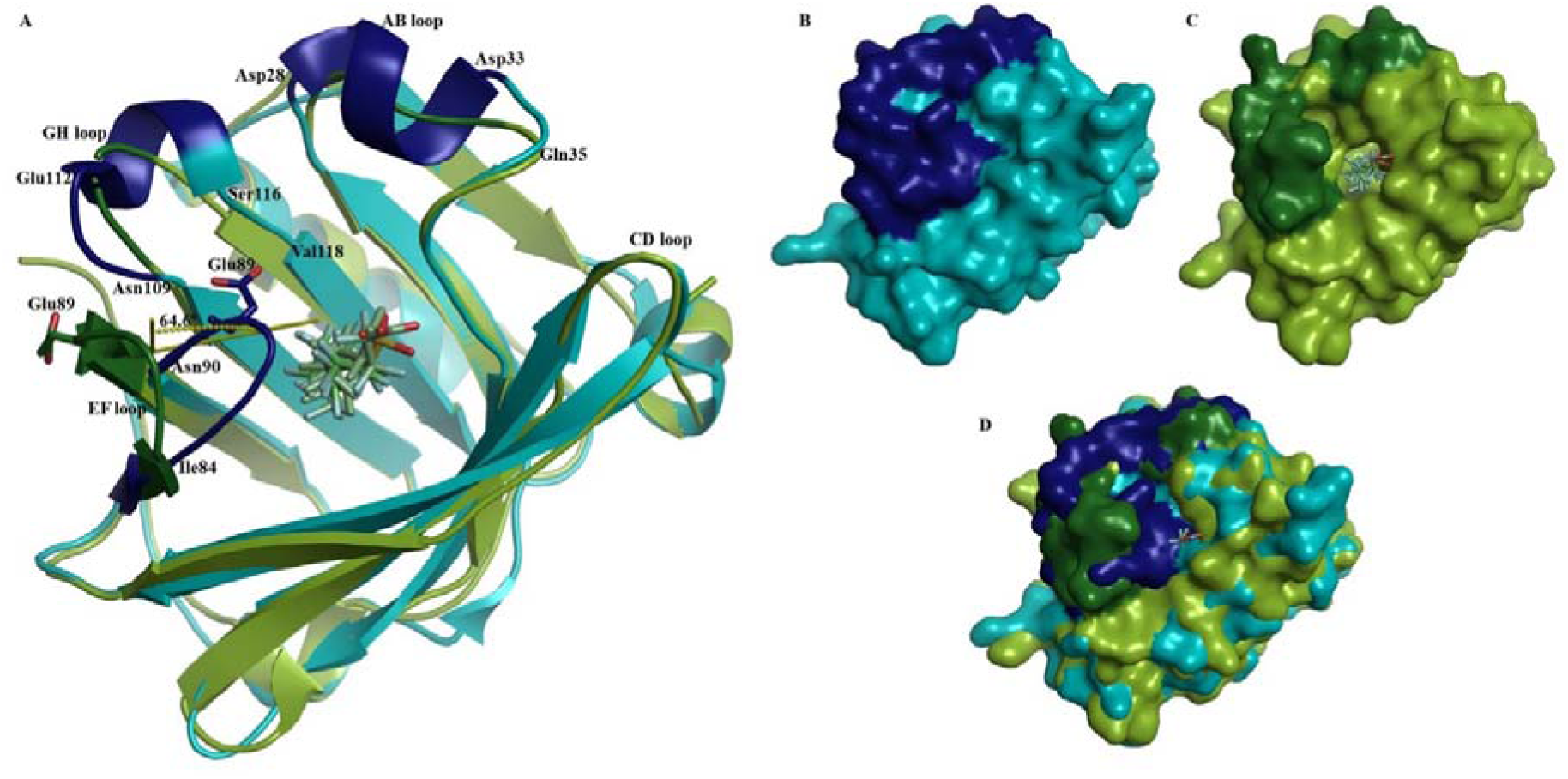
Comparative structural analysis of closed and open conformations of the β-lactoglobulin-PFAS complexes. A) Cartoon representation of superimposed structures of apo-β-lactoglobulin closed (blue) conformation (PDB ID: 2AKQ at pH 5.2, previous study) over open conformation (green) of β-lactoglobulin-PFAS complexes (present study), representing structural changes highlighted in dark green colour for open conformation and dark blue colour for closed conformation. B) Surface view of the apo-β-lactoglobulin closed (blue) conformation (PDB ID: 2AKQ, at pH 5.2, previous study) C) Surface view of open conformation (green) of β-lactoglobulin-PFAS complexes D) Superimposed surface view of closed (blue) and open (green) conformations of β-lactoglobulin.

Furthermore, specific changes in conformations of the AB and GH loops were also observed at pH 5.2. The residues Asp28 to Asp33 in the AB loop formed a small 3_10_ helix in the apo-β-lactoglobulin (at pH 5.2), while in β-lactoglobulin PFAS complexes and in apo-β-lactoglobulin at pH 6.2, 7.1 (PDB ID: 1BSY), and 8.2, these residues were present in loop conformation. Further, Glu112 to Ser116 of the GH loop were in helical conformation in the apo-form at pH 5.2, while were in loop conformation in apo-β-lactoglobulin at pH 6.2, 7.1 and 8.2 and in β-lactoglobulin PFAS complexes at pH 8.2, residues Ala111-Glu114 were missing and no helical conformation was observed in this loop (Figure 4A).^36, 52, 53, 35^

These structural changes in β-lactoglobulin in the presence of PFAS compounds (along with pH as an additional factor) provide insight into the PFAS binding mechanism of β-lactoglobulin. The formation of the complex with PFAS occupying the canonical retinol/fatty acid hydrophobic binding pocket potentially begins to speak to its ability to transport PFAS in PFAS-contaminated milk. Physiologically, this phenomenon could potentially increase PFAS concentration not only in neonates but also in children and adults.

### Molecular dynamics simulation analysis

Molecular dynamics simulation of crystal structures of β-lactoglobulin-PFDA, β-lactoglobulin-PFOS, β-lactoglobulin-PFOA complexes, and apo-β-lactoglobulin was performed for a duration of 100 ns in two independent runs for each system to study the PFAS binding dynamics of β-lactoglobulin. The MD simulation trajectory analysis revealed consistent binding of the PFAS molecule in the central calyx cavity of the β-lactoglobulin. The RMSD versus trajectory time plots showed convergence of all the trajectories after 20 ns within a narrow range of ∼ 0.20 to 0.30 nm with comparatively higher variations in the β-lactoglobulin-PFOS complex attributed to the presence of the sulphonic acid moiety (Figure 5A).^49^ The Rg versus time plots revealed consistent compactness of protein in all complexes within the range of 1.50 to 1.55 nm. The higher Rg values of the β-lactoglobulin-PFOS complex corresponding to the RMSD variations were observed, indicating significant conformational changes in the protein structure, conveying a minor reduction in the compactness of the protein (Figure 5B).^54^

**Figure 5:**
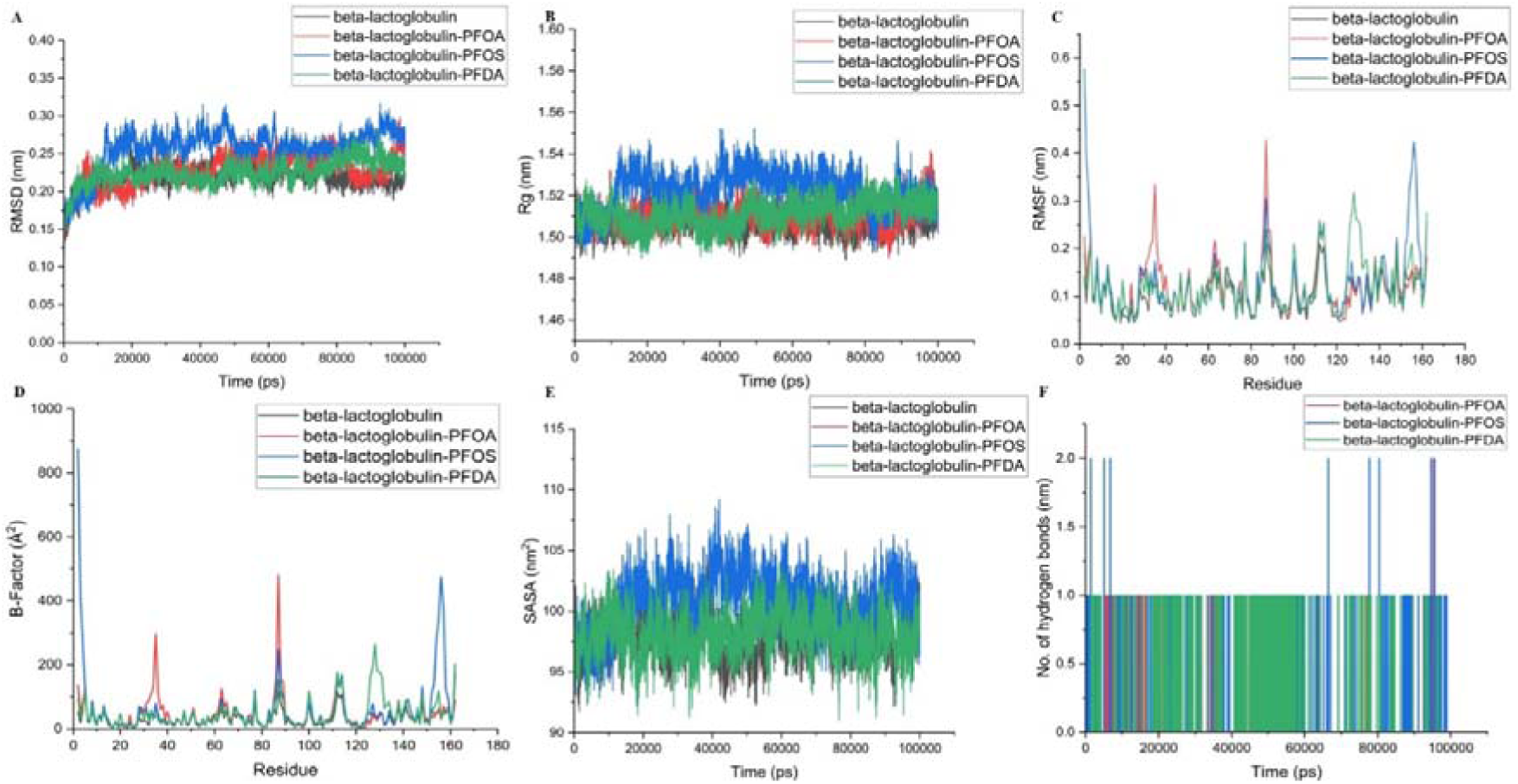
MD simulation analysis of apo-β-lactoglobulin and crystal structures of beta-lactoglobulin in complex with PFOA, PFOS, and PFDA. A) Root Mean Square Deviation (RMSD) versus time plots B) Radius of gyration (Rg) versus time plots C) Root Mean Square Fluctuations (RMSF) versus residue plots D) B-Factor versus residue plots during 100 ns of simulation trajectory E) Solvent Accessible Surface Area (SASA) versus time plots. F) Number of Hydrogen bonds versus time plots.

The RMSF versus residue plots showed significant fluctuations in the residues ranging from 80-85 and 110-120 in all three complexes in both replicates, which correspond to the opening of the EF loop and GH loop, respectively (Figure 5C and S1). Specific large fluctuations in residues 150-160 were observed in β-lactoglobulin-PFOS complex in both the replicates while the fluctuations in residues 30-40 and residues 127-133 of β-lactoglobulin-PFOA and PFDA respectively were observed in one of the two replicates indicates transient conformation sampling or moderate flexibility (Figure 5C and S1A). Similar variations in the B-factor versus residue plots were observed, indicating increased flexibility of the residues (Figure 5D and S1B). The 30-40 residue region, which displayed a high B-factor in β-lactoglobulin-PFOA complex in one of the replicates, indicates comparative flexibility of the AB loop in this complex. The 127-133 residue regions corresponding to the loop between the 8^th^ strand and the 3-turn alpha-helix lying at the base of the calyx showed high transient flexibility in the B-factor plot and thus high fluctuations in this region of the β-lactoglobulin-PFDA complex (Figure 5D). Such fluctuations at the base of the calyx were due to the comparatively deep insertion of the longer chain of PFDA into the central cavity of the calyx in one of the replicates, which conveyed conformational changes (Figure 5D). Further, the 152-158 residue region of the β-lactoglobulin-PFOS complex indicates the C-terminal small helix that experienced consistent effective fluctuations during the trajectories of both replicates (Figure 5D, S1B and 6).^33^ Thus the PFAS-specific alterations in B-factors suggest further investigation into the impact of PFAS chain-length, branching, and head-group on protein structures (side-chains, helices, sheets, loops, turns and binding cavities).

The solvent accessible surface area versus time plots shows relative increase in the surface area of protein in the β-lactoglobulin-PFOS complex compared to the other two complex structures of PFOA and PFDA with β-lactoglobulin due to the presence of bulky sulphonic acid polar group in PFOS compared to the carboxyl group in PFOA and PFDA. The overall changes in surface area fluctuated in the fine range of 92 to 107 nm^2^ in all three complex structures, indicating stable binding of PFAS compounds in the central cavity along with considerable changes in the protein structure without communicating any misfolding or unfolding (Figure 5E).^55^ The hydrogen bond versus trajectory time plots revealed consistent hydrogen bonding of PFAS compounds with the β-lactoglobulin. Trajectory analysis revealed the consistent hydrogen bond interactions of charged headgroups in PFAS compounds with Lys60 and/or Lys69 during the simulation trajectory (Figure 5F). Superimposition of trajectory frames reveals the allosteric structural changes at the base of the central calyx of β-lactoglobulin communicated by the binding of the PFAS molecules. The binding of PFDA in the central calyx, due to its longer hydrophobic chain tail compared to PFOA and PFOS, which further penetrated inside the cavity, resulted in an inward motion of the loop between Lue133 an Glu127 (Figure 6). Furthermore, in the complex structure of β-lactoglobulin-PFOS, the transition of helix to loop was discrete in the region ranging from Asn152 to Glu158, in comparison to the other two complexes during the trajectory of MD simulation.

**Figure 6:**
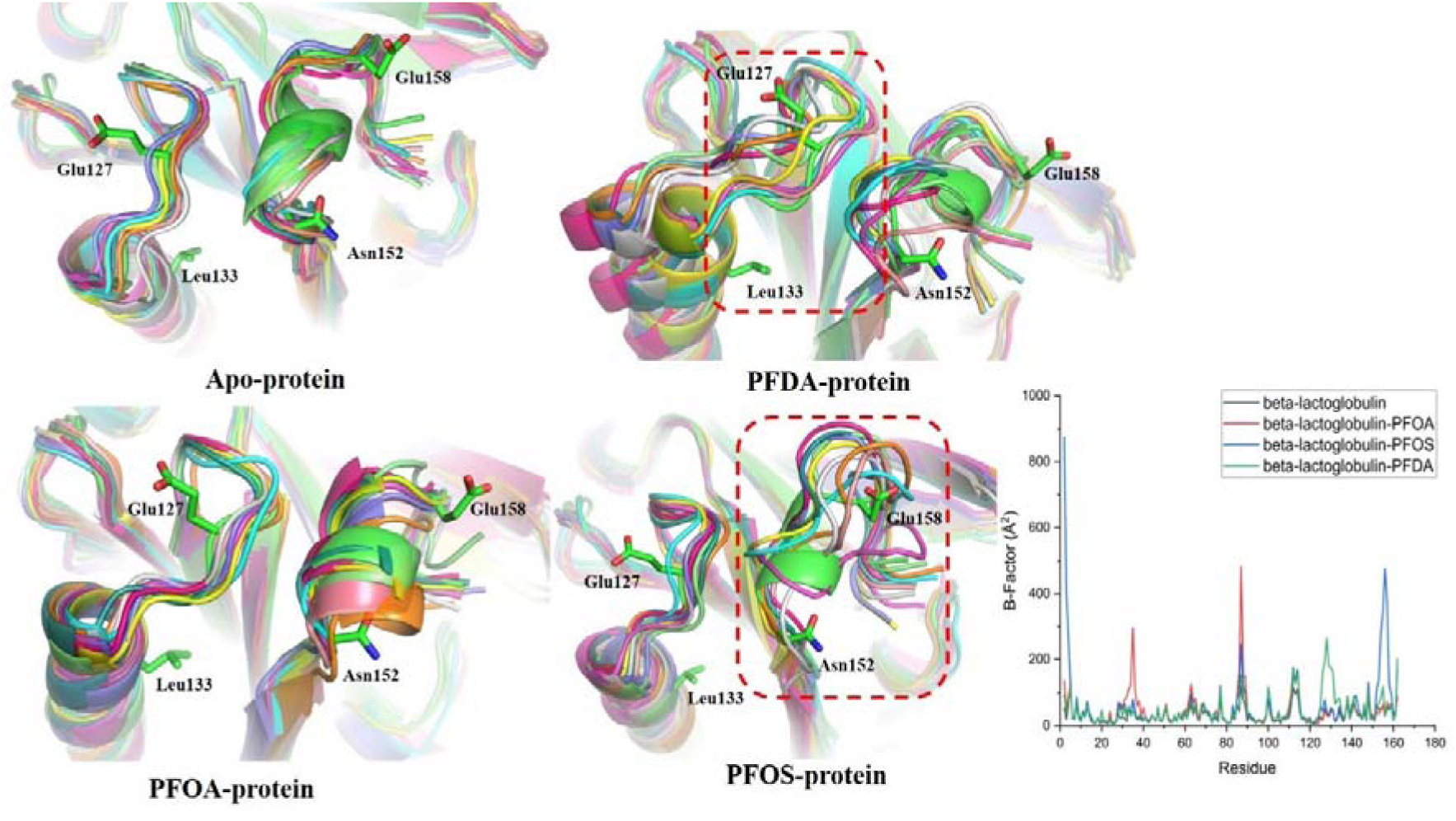
Superimposition of extracted frames at every 10 ns of the MD simulation trajectories of apo- β -lactoglobulin and crystal structures of β-lactoglobulin in complex with PFOA, PFOS, and PFDA. The regions showing major changes in the B-factor are highlighted with dashed frames. Structures in green colour show the initial conformations at 0 ns.

The secondary structure analysis elucidated some PFAS-induced variations in the coil, bend, and helical conformations in complex forms compared to the apo-β-lactoglobulin. One significant change was a major reduction in 3_10_-helix content observed in the β-lactoglobulin-PFDA complex during the simulation trajectory (Figure 7).^34^ 3_10_-helix acts as an intermediate for the folding/unfolding of α-helices.^34^ The major reduction of 3_10_-helix was observed in β-lactoglobulin-PFDA complex, which can be attributed to the long hydrophobic chain, which, upon binding into the central calyx, conveys major changes in the cavity, thereby changing the cavity volume (Figure 8).The number of residues contributing to other secondary structure elements that were analyzed mainly fluctuated, without showing consistent shifts (as expected in MD simulations at the 100 ns timescale performed in the present work). To further analyze structural conformations in three complexes, we performed cluster analysis, which revealed a significant number of structural conformations in the top 10 clusters of the complexes (Figure S2). The representative conformations of the top 10 clusters depicted consistent binding of the PFAS compounds in the central calyx of the β-lactoglobulin during the trajectory (Figure S2). Open extended conformations of the EF and GH loops were observed in all the complex clusters, while the AB loop was shifted inside the cavity. Significant changes in the cavity volume were evident in β-lactoglobulin-PFAS complexes during the trajectory. The volume of the cavity varied in the range of 487 Å^3^ to 1065 Å^3^ in β-lactoglobulin-PFDA complex, 498 Å^3^ to 1056 Å^3^ in β-lactoglobulin-PFOA complex, and 586 Å^3^ to 896 Å^3^ in β-lactoglobulin-PFOS complex with initial cavity volumes of 897Å^3^, 832 Å^3^ and 896 Å^3,^ respectively, for the three complexes (Figure 8).

**Figure 7:**
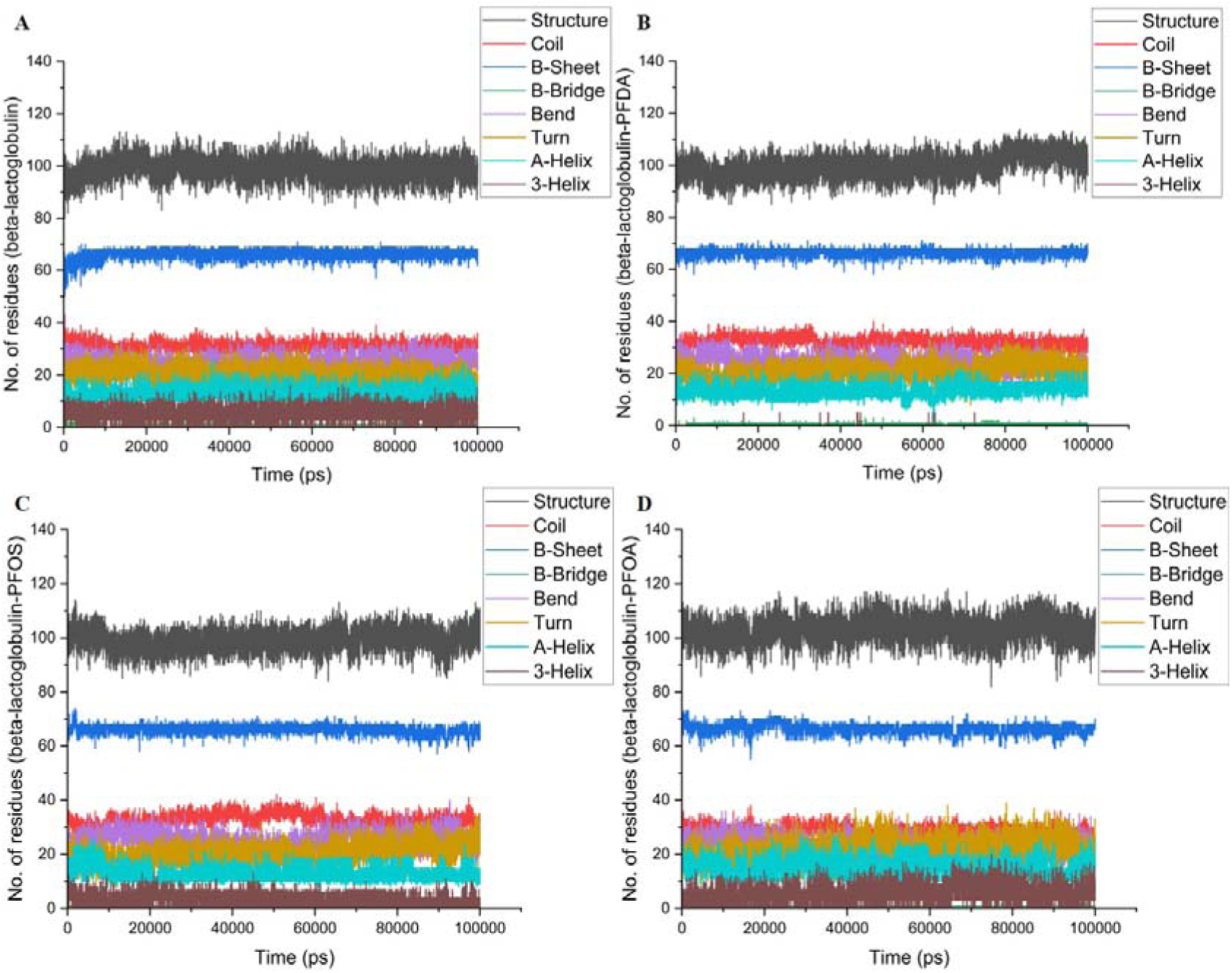
**MD simulation trajectory analysis for variation in secondary structures** in A) apo-β-lactoglobulin, B) β-lactoglobulin in complex with PFDA, C) β-lactoglobulin in complex with PFOS, D) β-lactoglobulin in complex with PFOA during the simulation trajectory of 100 ns. (Structure refers to number of residues contributing to all segments with defined secondary structure)

**Figure 8:**
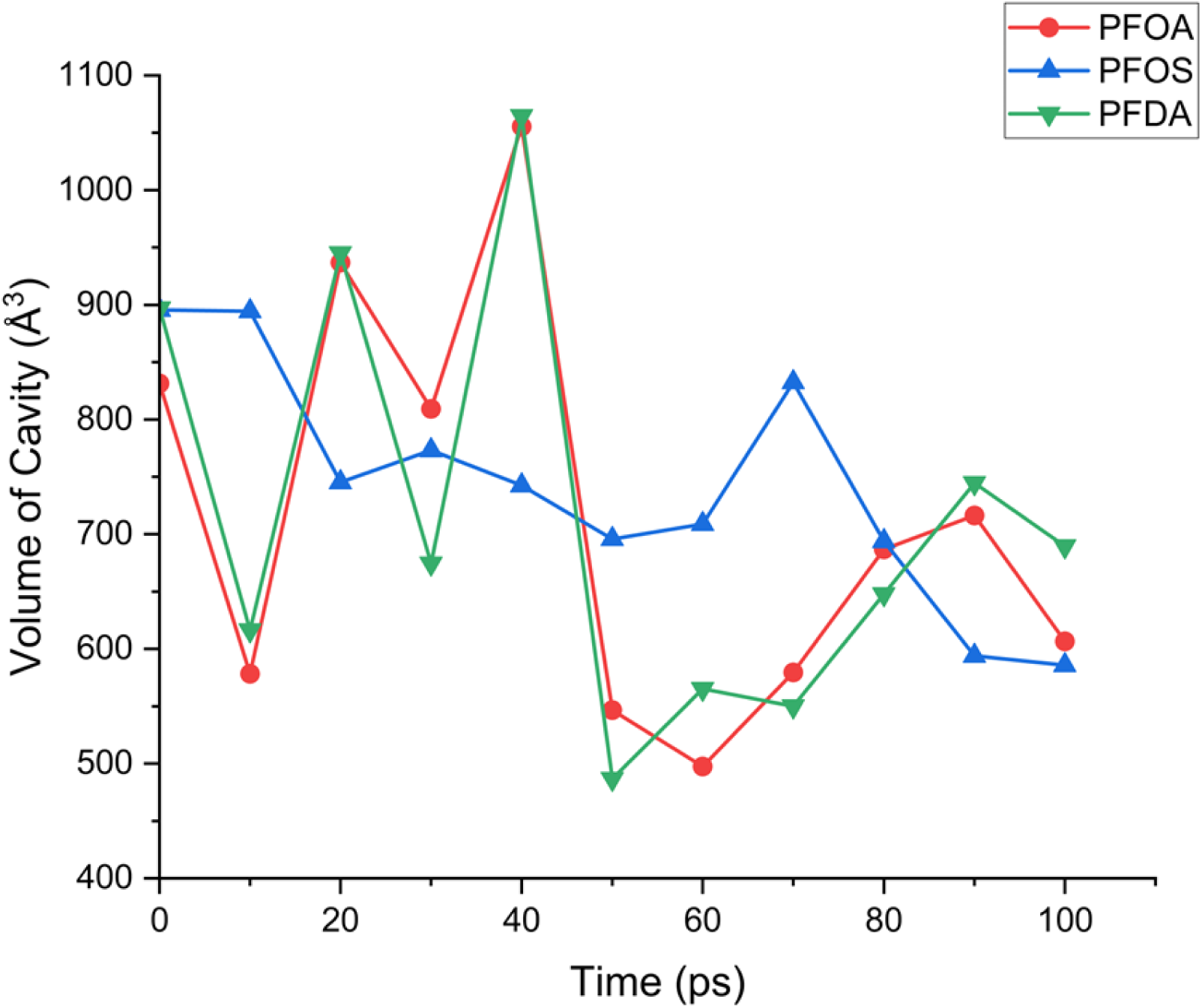
Line plot displaying the change in volume of central cavity (Å^3^) versus time of β-lactoglobulin in complex with PFOA, PFOS, and PFDA during the MD simulation trajectory of 100 ns.

MD simulation trajectories provide the means for estimating free energies of binding between PFAS molecules and β-lactoglobulin in three complexes. These free energies of binding and the individual components contributing to these free energies were estimated during the last 40 ns of two independently run simulation trajectories for each system. Total free energies of binding and several energy components, including van der Waals, electrostatic interaction energy, polar bond energy, and non-polar bond energy, had negative values, indicating favorable binding of the PFAS compounds in the central calyx of the β-lactoglobulin (Figure 9, S3). The total binding energies of PFDA, PFOS, and PFOA in the β-lactoglobulin calyx were -25 +/- 1.66, -21 +/- 1.91, and -23 +/- 1.50 kcal/mol, respectively. Out of all the evaluated energy components, the van der Waals interaction energy component was the leading contributor to the total binding energy, accounting for -32 +/- 2.56, -28 +/- 3.28, and -32 +/- 2.78 kcal/mol, respectively, for β-lactoglobulin and PFDA, PFOS, and PFOA complexes, therefore confirming effective PFAS binding in β-lactoglobulin.^51^

**Figure 9:**
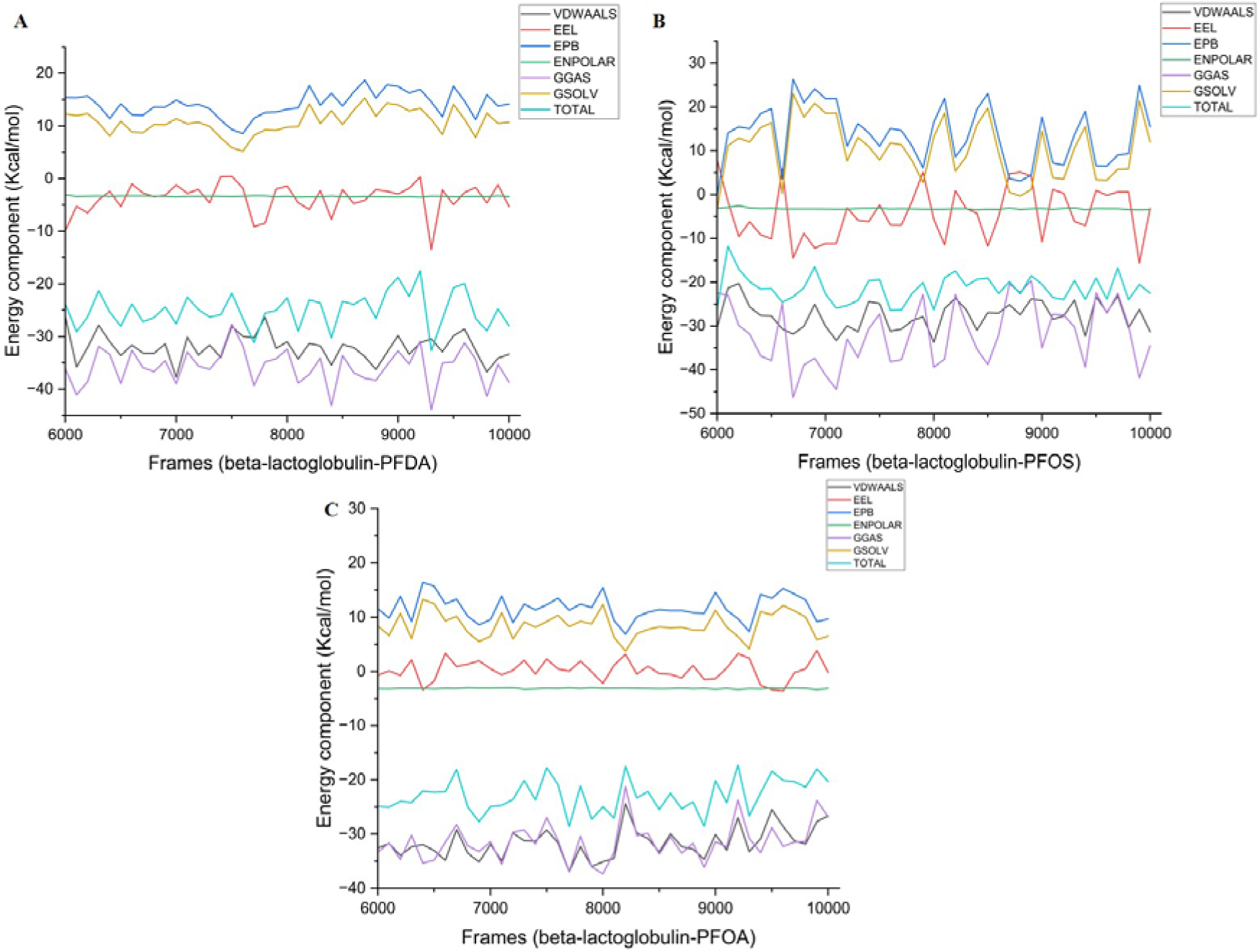
**End state free binding energy analysis** of beta-lactoglobulin in complex with A). PFDA, B) PFOS, C) PFOA showing variation in different energy components during the last 40 ns of the simulation trajectory. (VDWAALS: van der Waals, EEL: Electrostatic interaction energy, EPB: Polar bond energy, ENPOLAR: Non-polar bond energy, GGAS: VDWAALS + EEL, GSOLV: ENPOLAR + EPB, TOTAL = GGAS + GSOLV)

However, while several hypotheses associated with β-lactoglobulin PFAS binding have been offered, the present study provides the exact binding mechanism of PFAS within the central calyx cavity of the β-lactoglobulin by determining the crystal structures of β-lactoglobulin-PFDA, β-lactoglobulin-PFOS, and β-lactoglobulin-PFOA complexes and in addition, by analyzing their binding dynamics using an in-silico approach. The data also reveal that binding effects to biological assemblies (in this case a model globular protein) are also predicated upon the nature of the PFAS molecule. Features such as chain-length (which affects hydrophobicity), branching, nature of the head-group and its pKa are all likely to impact their interactions with biological systems from simple proteins to receptors, sub-cellular systems, matrices, tissues and organs.

Nevertheless, the findings divulged in the present analysis begin to provide valuable insights associated with PFAS transport to neonates via mother’s milk, cow milk or infant formulations, resulting in increased toxicity. The data can be beneficial in developing strategies to reduce toxicity by developing mechanisms that reduce PFAS contamination and by inhibiting PFAS binding in the central calyx of the milk protein β-lactoglobulin.

## Conclusion

This structural study identifies with atomic- and molecular-precision the interactions between three widely-prevalent PFAS molecules and β-lactoglobulin. It helps begin to reduce knowledge gaps in the first and subsequent molecular events that potentially onset PFAS-associated perturbations in β-lactoglobulin, which can then drive specific toxicological effects in humans. The work forms the foundation for cataloguing energetics, sterics, and intermolecular-interactions-based descriptors that are dependent upon PFAS head-group, chain length, and hydrophobicity as a function of protein side-chain and topology.

## Supporting information

Supporting Information Figure S1: MD simulation RMSF and B-factor plots (Duplicate run). Figure S2: Clustering analysis of the MD simulation trajecto

## Statements and Declarations

## Funding and Facilities

This work was supported by a DBT-funded projects entitled Translational and Structural Bioinformatics – BIC at the Department of Biotechnology, Indian Institute of Technology, Roorkee (BT/PR40141/BTIS/137/16/2021) and National Network Project of Department of Biotechnology, Indian Institute of Technology, Roorkee (BT/PR40142/BTIS/137/72/2023). MN is grateful to the National Institute of General Medical Sciences of the National Institutes of Health (NIH/NIGMS) under Award Number 1R16GM145575-01 for supporting this work.

## Authors Contributions

“The study conception and design were carried out by Pravindra Kumar and Mahesh Narayan. Data collection and analysis were performed by Shalja Verma, Anika Singh, Randhal S. Ramirez Orozco, and Lela Vukovic. The first draft of the manuscript was written by Shalja Verma, Pravindra Kumar, and Mahesh Narayan. All authors commented on previous versions of the manuscript and read and approved the final manuscript.”

## Ethics approval

Not applicable

## Declaration of generative AI

AI-based tool has not been used for this manuscript

## Accession codes

The crystal structures of PFDA-β-lactoglobulin complex (9WNW), PFOA-β-lactoglobulin complex (9WNV), and PFOS-β-lactoglobulin complex (9WNX) can be accessed via the Protein Data Bank. The UniProt ID of β-lactoglobulin is P02754

## Supporting Information

Figure S1: MD simulation RMSF and B-factor plots (Duplicate run).

Figure S2: Clustering analysis of the MD simulation trajectories.

Figure S3: End state free binding energy analysis plots (Duplicate run).

## For Table of Contents Only

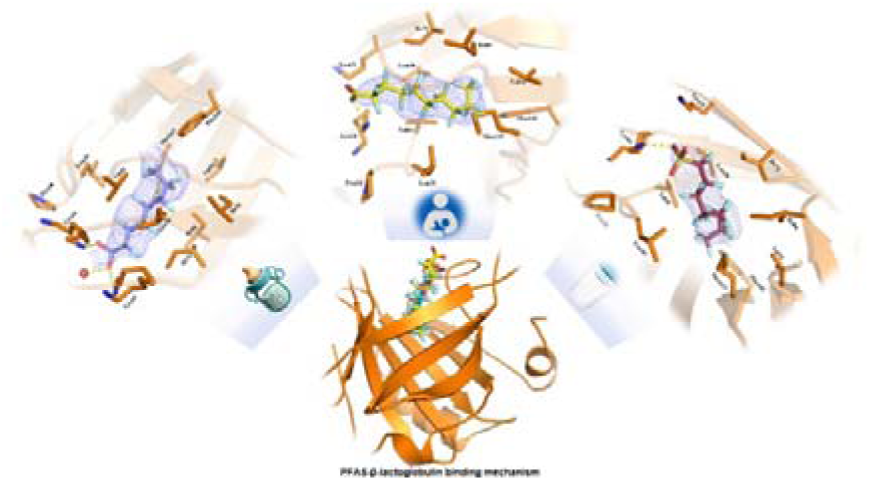

