## Supporting Information Figure S1: MD simulation RMSF and B-factor plots (Duplicate run). Figure S2: Clustering analysis of the MD simulation trajecto for "Structural insights into PFAS-β-lactoglobulin binding mechanism mediating PFAS toxicity"

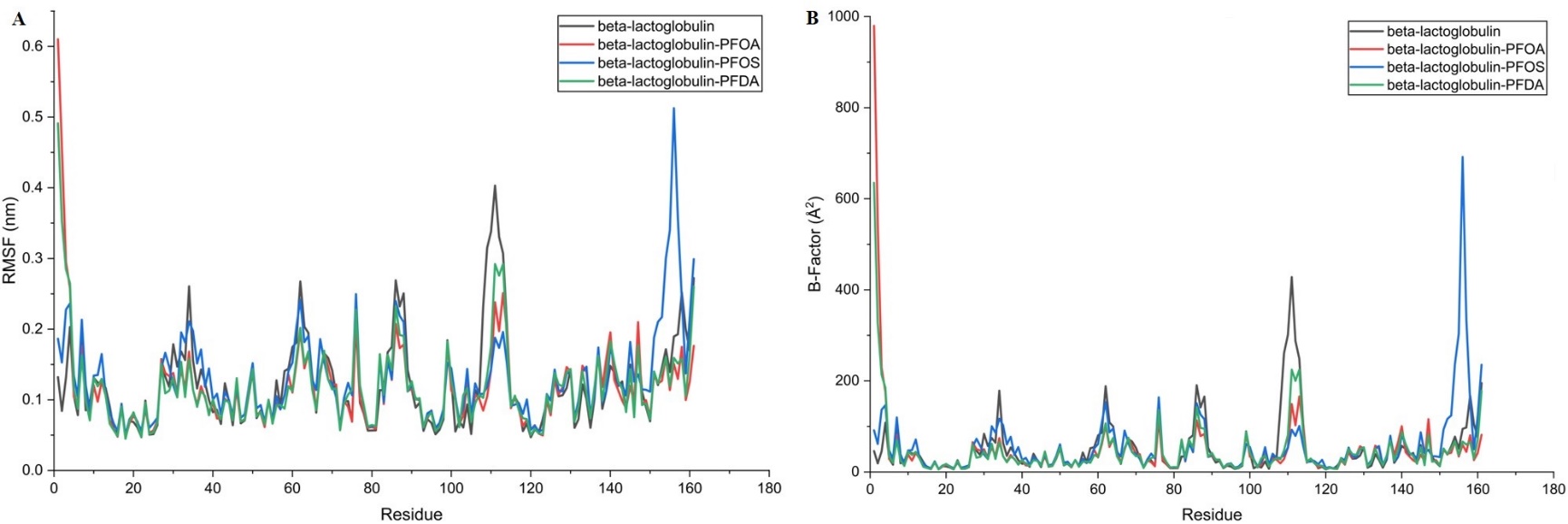

**Figure S1:** **MD simulation analysis (Duplicate run) of apo-β-lactoglobulin and crystal structures of beta-lactoglobulin in complex with PFOA, PFOS, and PFDA.** A) Root Mean Square Fluctuations (RMSF) versus residue plots B) B-Factor versus residue plots during 100 ns of simulation trajectory

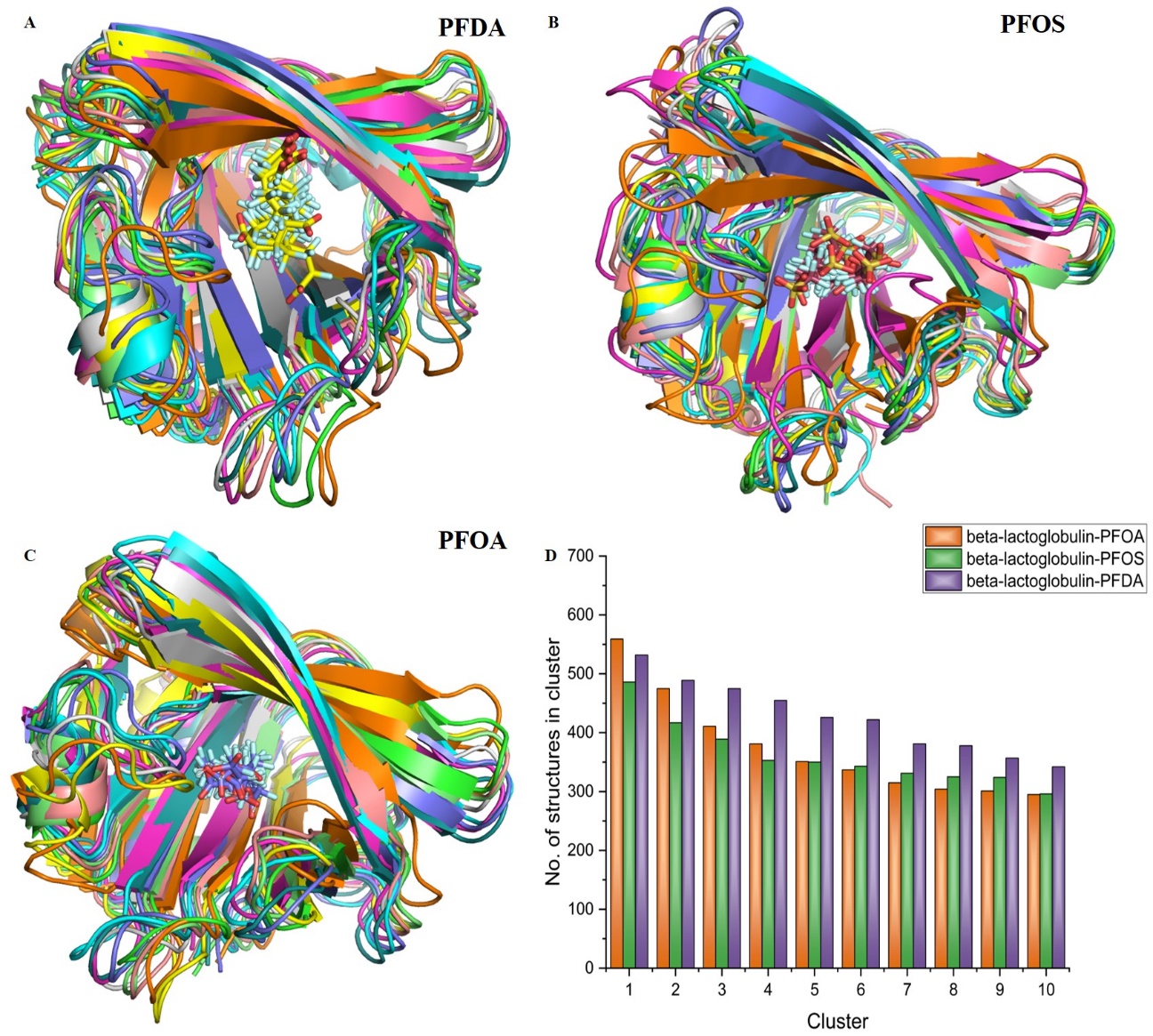

**Figure S2: Clustering analysis of the MD simulation trajectories.** A) Superimposition of representative conformations of the top 10 clusters analyzed from the simulation trajectory of beta-lactoglobulin-PFDA complex B) Superimposition of representative conformations of the top 10 clusters analyzed from the simulation trajectory of beta-lactoglobulin-PFOS complex C) Superimposition of representative conformations of the top 10 clusters analyzed from the simulation trajectory of beta-lactoglobulin-PFOA complex D) Bar plot showing number of structures identified in top 10 cluster from the simulation trajectories of beta-lactoglobulin-PFASs complexes.

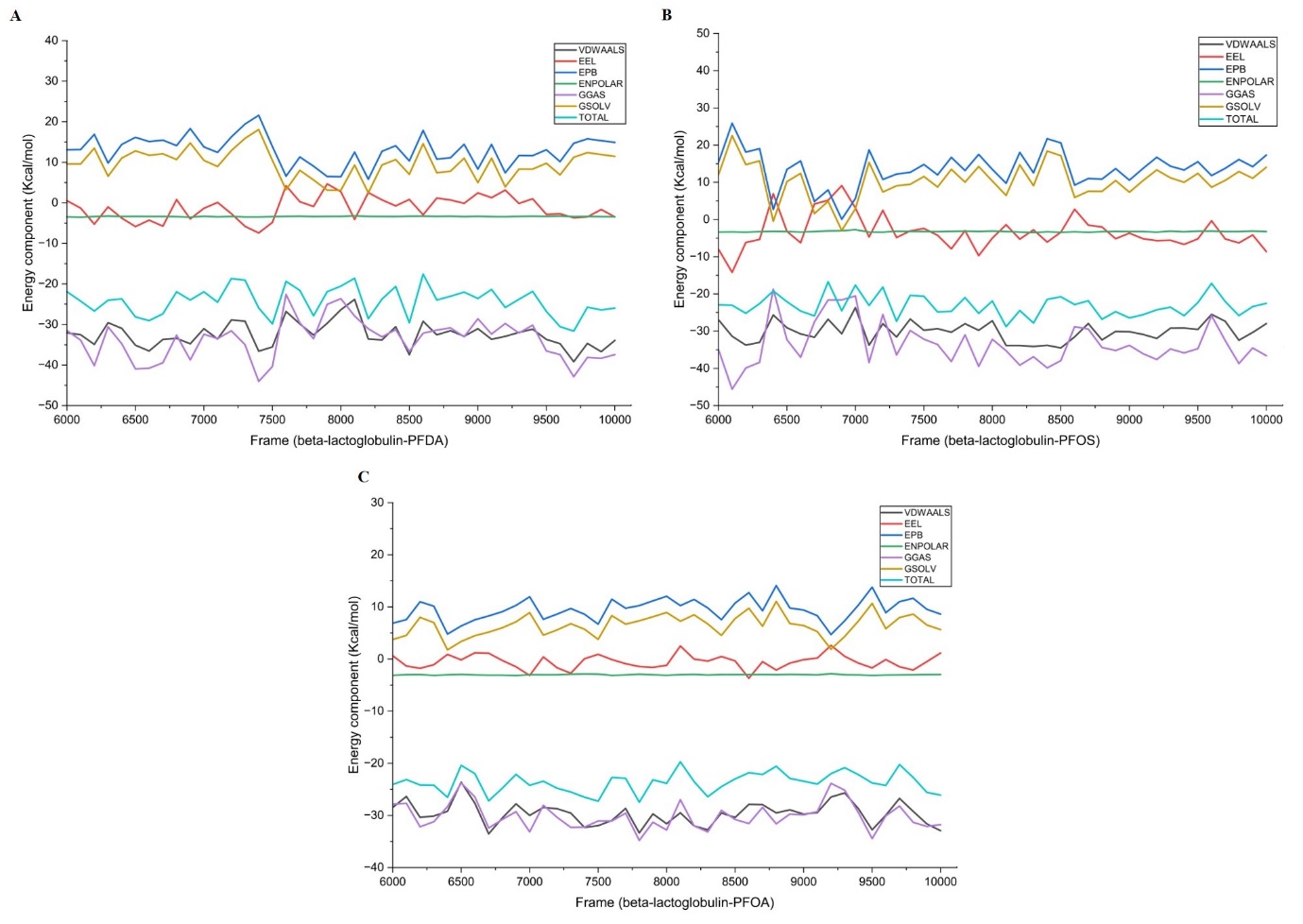

**Figure S3: End state free binding energy analysis (Duplicate run)** of beta-lactoglobulin in complex with A). PFDA, B) PFOS, C) PFOA showing variation in different energy components during the last 40 ns of the simulation trajectory. (VDWAALS: van der Waals, EEL: Electrostatic interaction energy, EPB: Polar bond energy, ENPOLAR: Non-polar bond energy, GGAS: VDWAALS + EEL, GSOLV: ENPOLAR + EPB, TOTAL = GGAS + GSOLV)
